## supplemental data for "Identification and copy number variant analysis of enhancer regions of genes causing spinocerebellar ataxia"

**Supplementary table 1**. Promoter viewpoints and 4C-templates used for each viewpoint of *ATXN1*, *ATXN3*, *TBP* and *ITPR1*.

| Gene | Promoter viewpoint (Hg19) | 4C-template |
|---|---|---|
| *ATXN1* | Chr6:16761086-16761915 | *NlaIII/Csp6I* |
| *ATXN3* | Chr14:92572894-92573491 | *NlaIII/DpnII* |
| *TBP* | Chr6:170863371-170863896 | *NlaIII/DpnII* |
| *ITPR1* | Chr3:4533821-4535043 | *NlaIII/DpnII* |

**Supplementary table 2**. Reading, non-reading and universal primers carrying the Illumina adaptors for amplification of regions interacting with *ATXN1*, *ATXN3*, *TBP* and *ITPR1* promoter viewpoints**.** Reading and non-reading primers (in bold) carry an additional 5' tail (red and green respectively) which anneal to the 3' end of the forward-universal primer (in red) and index-reverse primers (in green). Three index-reverse primers with different barcodes (yellow) are used for the three replicates of each promoter viewpoint.

| Viewpoint | Primer | Sequence (5' → 3') |
|---|---|---|
| *ATXN1* promoter | Reading | TACACGACGCTCTTCCGATCT–GA–**TCAAGGATCCGACTTCATG** |
| | Non-reading | ACTGGAGTTCAGACGTGTGCTCTTCCGATCT-**TTTCTACTACAGTGGCGGAG** |
| | Forward-universal | AATGATACGGCGACCACCGAGATCTACACTCTTTCCCTACACGACGCTCTTCCGATCT |
| | Index-reverse | CAAGCAGAAGACGGCATACGAGATTGTTGACTGTGACTGGAGTTCAGACGTGTGCT |
| | | CAAGCAGAAGACGGCATACGAGATACGGAACTGTGACTGGAGTTCAGACGTGTGCT |
| | | CAAGCAGAAGACGGCATACGAGATTCTGACATGTGACTGGAGTTCAGACGTGTGCT |
| *ATXN3* promoter | Reading | TACACGACGCTCTTCCGATCT–ACG–**TCCAGACAAATAAACATG** |
| | Non-reading | ACTGGAGTTCAGACGTGTGCTCTTCCGATCT-**CTCCTAGCTTCTGTTAGC** |
| | Forward-universal | AATGATACGGCGACCACCGAGATCTACACTCTTTCCCTACACGACGCTCTTCCGATCT |
| | Index-reverse | CAAGCAGAAGACGGCATACGAGATGATCTGGTGACTGGAGTTCAGACGTGTGCT |
| | | CAAGCAGAAGACGGCATACGAGATTCAAGTGTGACTGGAGTTCAGACGTGTGCT |

| | | |
|---|---|---|
| | | CAAGCAGAAGACGGCATACGAGATCTGATCGTGACTGGAGTTCAGACGTGTGCT |
| *TBP* promoter | Reading | TACACGACGCTCTTCCGATCT-CTAGG-**ATCCGCTTCTTCTCCATG** |
| | Non-reading | ACTGGAGTTCAGACGTGTGCTCTTCCGATCT-**CCCGAACCGTCAGAAAAGAG** |
| | Forward-universal | AATGATACGGCGACCACCGAGATCTACACTCTTTCCCTACACGACGCTCTTCCGATCT |
| | Index-reverse | CAAGCAGAAGACGGCATACGAGATCGTGATGTGACTGGAGTTCAGACGTGTGCT |
| | | CAAGCAGAAGACGGCATACGAGATACATCGGTGACTGGAGTTCAGACGTGTGCT |
| | | CAAGCAGAAGACGGCATACGAGATGCCTAAGTGACTGGAGTTCAGACGTGTGCT |
| *ITPR1* promoter | Reading | TACACGACGCTCTTCCGATCT–G–**TTTAAAGTGCAGTAACCATG** |
| | Non-reading | ACTGGAGTTCAGACGTGTGCTCTTCCGATCT-**CCAAACGCATTATAGCATCC** |
| | Forward-universal | AATGATACGGCGACCACCGAGATCTACACTCTTTCCCTACACGACGCTCTTCCGATCT |
| | Index-reverse | CAAGCAGAAGACGGCATACGAGATTGGTCAGTGACTGGAGTTCAGACGTGTGCT |
| | | CAAGCAGAAGACGGCATACGAGATCACTGTGTGACTGGAGTTCAGACGTGTGCT |
| | | CAAGCAGAAGACGGCATACGAGATATTGGCGTGACTGGAGTTCAGACGTGTGCT |

**Supplementary table 3**. First PCR condition for the 4C sequencing library preparation.

| **Component** | $\mu$l |
|---|---|
| Buffer 1 | 2.5 |
| dNTP (10 mM) | 0.5 |
| Reading primer (10 µM) | 3.5 |
| Non-reading primer (10 µM) | 3.5 |
| Expand long template polymerase | 0.35 |
| Template (150 ng) | x |
| MilliQ | x |
| **Total volume** | **25** |

**Supplementary table 4.** Program of the first PCR for the 4C sequencing library preparation.

<table>
<tr><th>Temp (°C)</th><th>Time</th><th>Cycles</th></tr>
<tr><td>94</td><td>2 min</td><td></td></tr>
<tr><td>94</td><td>15 sec</td><td rowspan="3">16</td></tr>
<tr><td>54</td><td>1 min</td></tr>
<tr><td>68</td><td>3 min</td></tr>
<tr><td>68</td><td>7 min</td><td></td></tr>
<tr><td>10</td><td>∞</td><td></td></tr>
</table>

**Supplementary table 5.** Second PCR condition for the 4C sequencing library preparation.

| **Component** | ***µ*l** |
|---|---|
| Buffer 1 | 2.5 |
| dNTP (10 mM) | 0.5 |
| Primer mix (Forward-universal and index-reverse (5 µM)) | 2.5 |
| Expand long template polymerase | 0.35 |
| First round PCR product | 2.5 |
| MilliQ | x |
| **Total volume** | **25** |

**Supplementary table 6**. Program of the second PCR for the 4C sequencing library preparation.

| **Temp (°C)** | **Time** | **Cycles** |
|---|---|---|
| 94 | 2 min | |
| 94 | 15 sec | 20 |
| 60 | 1min | |
| 68 | 3 min | |
| 68 | 7 min | |
| 10 | ∞ | |

**Supplementary table 7.** PutE viewpoints and the 4C-template used for each viewpoint.

| **PutE** | **PutE viewpoint (Hg19)** | **4C-template** |
|---|---|---|
| *ATXN1*-PutE1 | chr6:16321036-16321612 | *NlaIII*/*DpnII* |
| *ATXN1*-PutE2 | chr6:16803517-16804414 | *NlaIII*/*Csp6I* |
| *ATXN3*-PutE1 | chr14:91975457-91975776 | *NlaIII*/*DpnII* |
| *ATXN3*-PutE2 | chr14:92505411-92506375 | *NlaIII*/*Csp6I* |
| *ATXN3*-PutE3 | chr14:93062668-93063126 | *NlaIII*/*Csp6I* |
| *ATXN3*-PutE4 | chr14:93215111-93216202 | *NlaIII*/*Csp6I* |
| *ATXN3*-PutE5 | chr14:93257696-93258172 | *NlaIII*/*DpnII* |
| *TBP*-PutE1 | chr6:170150992-170151453 | *NlaIII*/*DpnII* |
| *TBP*-PutE2 | chr6:170190125-170190807 | *NlaIII*/*DpnII* |
| *ITPR1*-PutE1 | chr3:3839485-3839753 | *NlaIII*/*DpnII* |
| *ITPR1*-PutE2 | chr3:4340769-4341564 | *NlaIII*/*DpnII* |
| *ITPR1*-PutE3 | chr3:4511190-4511674 | *NlaIII*/*DpnII* |
| *ITPR1*-PutE4 | chr3:4762537-4762939 | *NlaIII*/*Csp6I* |
| *ITPR1*-PutE5 | chr3:4790941-4791414 | *NlaIII*/*DpnII* |
| *ITPR1*-PutE6 | chr3:4908651-4909056 | *NlaIII*/*DpnII* |
| *ITPR1*-PutE7 | chr3:5020371-5020891 | *NlaIII*/*DpnII* |

**Supplementary table 8**. Reading, non-reading and universal primers carrying the Illumina adaptors for amplification of regions interacting with the PutE viewpoints. Reading and non-reading primers (in bold) carry an addition 5' tail (red and green, respectively) that anneals to the 3' end of the forward-universal (in red) and index-reverse primers (in green). Two index-reverse primers with different barcodes (yellow) were used for the two biological replicates of each PutE viewpoint.

| PutE | Primer | Sequence (5' → 3') |
|---|---|---|
| *ATXN1*-PutE1 | Reading | TACACGACGCTCTTCCGATCT–TA–**CTCAACCAGAATCAAGGCATG** |
| | Non-reading | ACTGGAGTTCAGACGTGTGCTCTTCCGATCT-**AGTAGGTTAGCTGCTTGGA** |
| | Forward-universal | AATGATACGGCGACCACCGAGATCTACACTCTTTCCCTACACGACGCTCTTCCGATCT |
| | Index-reverse | CAAGCAGAAGACGGCATACGAGATGCCTAAGTGACTGGAGTTCAGACGTGTGCT |
| | | CAAGCAGAAGACGGCATACGAGATTGGTCAGTGACTGGAGTTCAGACGTGTGCT |
| *ATXN1*-PutE2 | Reading | TACACGACGCTCTTCCGATCT-A-**GGTTTTTCTAAGAAGGCATG** |
| | Non-reading | ACTGGAGTTCAGACGTGTGCTCTTCCGATCT-**CTTGGCTGTTACAATTCATC** |
| | Forward-universal | AATGATACGGCGACCACCGAGATCTACACTCTTTCCCTACACGACGCTCTTCCGATCT |
| | Index-reverse | CAAGCAGAAGACGGCATACGAGATAAGGCCACGTGACTGGAGTTCAGACGTGTGCT |
| | | CAAGCAGAAGACGGCATACGAGATTCCGAAACGTGACTGGAGTTCAGACGTGTGCT |
| *ATXN3*-PutE1 | Reading | TACACGACGCTCTT[illegible]-**CGTGGAGCGAGCGCCATG** |
| | Non-reading | ACTGGAGTTCAGAC[illegible]TCCGATCT-**AGTGTGGAGAGGTCGAGGA** |
| | Forward-universal | AATGATACGGCGACCACCGAGATCTACACTCTTTCCCTACACGACGCTCTTCCGATCT |

| | | |
|---|---|---|
| | Index-reverse | CAAGCAGAAGACGGCATACGAGATTGTTGACTGTGACTGGAGTTCAGACGTGTGCT |
| | | CAAGCAGAAGACGGCATACGAGATACGGAACTGTGACTGGAGTTCAGACGTGTGCT |
| *ATXN3*-PutE2 | Reading | TACACGACGCTCTTCCGATCT-ATAGG- **TTATGAAACACGGCCATG** |
| | Non-reading | ACTGGAGTTCAGACGTGTGCTCTTCCGATCT-**ACGTAGGAGCAGTTTCACTCT** |
| | Forward-universal | AATGATACGGCGACCACCGAGATCTACACTCTTTCCCTACACGACGCTCTTCCGATCT |
| | Index-reverse | CAAGCAGAAGACGGCATACGAGATCGTGATGTGACTGGAGTTCAGACGTGTGCT |
| | | CAAGCAGAAGACGGCATACGAGATACATCGGTGACTGGAGTTCAGACGTGTGCT |
| *ATXN3*-PutE3 | Reading | TACACGACGCTCTTCCGATCT-TGG-**TCTATGAATACCCACATG** |
| | Non-reading | ACTGGAGTTCAGACGTGTGCTCTTCCGATCT-**TTGGAATTACCCTTGGAGTC** |
| | Forward-universal | AATGATACGGCGACCACCGAGATCTACACTCTTTCCCTACACGACGCTCTTCCGATCT |
| | Index-reverse | CAAGCAGAAGACGGCATACGAGATCGGGACGGGTGACTGGAGTTCAGACGTGTGCT |
| | | CAAGCAGAAGACGGCATACGAGATGTGCGGACGTGACTGGAGTTCAGACGTGTGCT |
| *ATXN3*-PutE4 | Reading | TACACGACGCTCTTCCGATCT-ATCA-**ATCCATTCGAGTGCCATG** |
| | Non-reading | ACTGGAGTTCAGACGTGTGCTCTTCCGATCT-**ACAAGGTTGGCACACAGGT** |
| | Forward-universal | AATGATACGGCGACCACCGAGATCTACACTCTTTCCCTACACGACGCTCTTCCGATCT |

| | | |
|---|---|---|
| | Index-reverse | CAAGCAGAAGACGGCATACGAGATATCCACTCGTGACTGGAGTTCAGACGTGTGCT |
| | | CAAGCAGAAGACGGCATACGAGATATATCAGTGTGACTGGAGTTCAGACGTGTGCT |
| *ATXN3*-PutE5 | Reading | TACACGACGCTCTTCCGATCT-GA-**CTGTGACACGTTACCATG** |
| | Non-reading | ACTGGAGTTCAGACGTGTGCTCTTCCGATCT-**AAACTTCATCCTTAAGCTCT** |
| | Forward-universal | AATGATACGGCGACCACCGAGATCTACACTCTTTCCCTACACGACGCTCTTCCGATCT |
| | Index-reverse | CAAGCAGAAGACGGCATACGAGATCGGGACGGGTGACTGGAGTTCAGACGTGTGCT |
| | | CAAGCAGAAGACGGCATACGAGATGTGCGGACGTGACTGGAGTTCAGACGTGTGCT |
| *TBP*-PutE1 | Reading | TACACGACGCTCTTCCGATCT–TA-**TGCCTCCTTCTCGAACATG** |
| | Non-reading | ACTGGAGTTCAGACGTGTGCTCTTCCGATCT-**AGTCGATTGGAAGGGTGGTT** |
| | Forward universal | AATGATACGGCGACCACCGAGATCTACACTCTTTCCCTACACGACGCTCTTCCGATCT |
| | Index-reverse | CAAGCAGAAGACGGCATACGAGATGATCTGGTGACTGGAGTTCAGACGTGTGCT |
| | | CAAGCAGAAGACGGCATACGAGATTCAAGTGTGACTGGAGTTCAGACGTGTGCT |
| *TBP*-PutE2 | Reading | TACACGACGCTCTTCCGATCT–GCAC-**GTAGGAGTTCCGACGCATG** |
| | Non-reading | ACTGGAGTTCAGACGTGTGCTCTTCCGATCT-**CCTTTGCGAATCGATGGGGA** |
| | Forward-universal | AATGATACGGCGACCACCGAGATCTACACTCTTTCCCTACACGACGCTCTTCCGATCT |
| | Index-reverse | CAAGCAGAAGACGGCATACGAGATAAGCTAGTGACTGGAGTTCAGACGTGTGCT |

| | | |
|---|---|---|
| | | CAAGCAGAAGACGGCATACGAGATGTAGCCGTGACTGGAGTTCAGACGTGTGCT |
| *ITPR1*-PutE1 | Reading | TACACGACGCTCTTCCGATCT–CG–**TATCAGACAAACTCCATG** |
| | Non-reading | ACTGGAGTTCAGACGTGTGCTCTTCCGATCT-**CAGTCGGAGCAATTAATGAT** |
| | Forward-universal | AATGATACGGCGACCACCGAGATCTACACTCTTTCCCTACACGACGCTCTTCCGATCT |
| | Index-reverse | CAAGCAGAAGACGGCATACGAGATCACTGTGTGACTGGAGTTCAGACGTGTGCT |
| | | CAAGCAGAAGACGGCATACGAGATATTGGCGTGACTGGAGTTCAGACGTGTGCT |
| *ITPR1*-PutE2 | Reading | TACACGACGCTCTTCCGATCT-C-**TTAGCTATTATTTCCCATG** |
| | Non-reading | ACTGGAGTTCAGACGTGTGCTCTTCCGATCT-**GGTTTCACCGAATTGACC** |
| | Forward-universal | AATGATACGGCGACCACCGAGATCTACACTCTTTCCCTACACGACGCTCTTCCGATCT |
| | Index-reverse | CAAGCAGAAGACGGCATACGAGATAAGGCCACGTGACTGGAGTTCAGACGTGTGCT |
| | | CAAGCAGAAGACGGCATACGAGATTCCGAAACGTGACTGGAGTTCAGACGTGTGCT |
| *ITPR1*-PutE3 | Reading | TACACGACGCTCTTCCGATCT-G-**CCTAGGTCATACTTCATG** |
| | Non-reading | ACTGGAGTTCAGACGTGTGCTCTTCCGATCT-**GATGGAACTTTGTTGGTTCT** |
| | Forward-universal | AATGATACGGCGACCACCGAGATCTACACTCTTTCCCTACACGACGCTCTTCCGATCT |
| | Index-reverse | CAAGCAGAAGACGGCATACGAGATTACGTACGGTGACTGGAGTTCAGACGTGTGCT |

| | | |
|---|---|---|
| | | CAAGCAGAAGACGGCATACGAGATAAGCTAGTGACTGGAGTTCAGACGTGTGCT |
| *ITPR1*-PutE4 | Reading | TACACGACGCTCTTCCGATCT-A-**AAGTTGTATAATTTGGTCATG** |
| | Non-reading | ACTGGAGTTCAGACGTGTGCTCTTCCGATCT-**GCAGTGAGCCTAAATTCCA** |
| | Forward-universal | AATGATACGGCGACCACCGAGATCTACACTCTTTCCCTACACGACGCTCTTCCGATCT |
| | Index-reverse | CAAGCAGAAGACGGCATACGAGATGTAGCCGTGACTGGAGTTCAGACGTGTGCT |
| | | CAAGCAGAAGACGGCATACGAGATTACAAGGTGACTGGAGTTCAGACGTGTGCT |
| *ITPR1*-PutE5 | Reading | TACACGACGCTCTTCCGATCT–GC–**GCAACTGAAAAGGTGCCATG** |
| | Non-reading | ACTGGAGTTCAGACGTGTGCTCTTCCGATCT-**GGGCCTCCATATACACTGA** |
| | Forward-universal | AATGATACGGCGACCACCGAGATCTACACTCTTTCCCTACACGACGCTCTTCCGATCT |
| | Index-reverse | CAAGCAGAAGACGGCATACGAGATGATCTGGTGACTGGAGTTCAGACGTGTGCT |
| | | CAAGCAGAAGACGGCATACGAGATTCAAGTGTGACTGGAGTTCAGACGTGTGCT |
| *ITPR1*-PutE6 | Reading | TACACGACGCTCTTCCGATCT—**TTTTTGTGAGCCTTTCATG** |
| | Non-reading | ACTGGAGTTCAGACGTGTGCTCTTCCGATCT-**CAAACGGATTTTCTCTGAACTC** |
| | Forward-universal | AATGATACGGCGACCACCGAGATCTACACTCTTTCCCTACACGACGCTCTTCCGATCT |
| | Index-reverse | CAAGCAGAAGACGGCATACGAGATCTGATCGTGACTGGAGTTCAGACGTGTGCT |
| | | CAAGCAGAAGACGGCATACGAGATTGTTGACTGTGACTGGAGTTCAGACGTGTGCT |

| | | |
|---|---|---|
| *ITPR1*-PutE7 | Reading | TACACGACGCTCTTCCGATCT—**CAACACGTGAGACTCATG** |
| | Non-reading | ACTGGAGTTCAGACGTGTGCTCTTCCGATCT-**AGTTCCCGTCTCCTGGGA** |
| | Forward-universal | AATGATACGGCGACCACCGAGATCTACACTCTTTCCCTACACGACGCTCTTCCGATCT |
| | Index-reverse | CAAGCAGAAGACGGCATACGAGATACGGAACTGTGACTGGAGTTCAGACGTGTGCT |
| | | CAAGCAGAAGACGGCATACGAGATTCTGACATGTGACTGGAGTTCAGACGTGTGCT |

**Supplementary table 9.** Primers used to amplify R-PutEs of *ATXN1*, *ATXN3*, *TBP* and *ITPR1*, a previously reported validated enhancer (positive control) [12] and a random 1479 bp DNA fragment (negative control) for cloning into the pGL4.23 Firefly luciferase reporter vector. Primer sequences are shown in bold and the restriction sites in orange.

| R-PutEs | Cloned region (Hg19) | Primers Sequence (5' → 3') | |
|---|---|---|---|
| *ATXN1*-R-PutE1 | chr6:16319330-16322021 | Forward | AAAAGATATC**TCCCTGTTCCTAGGGAGGCTGAAG** |
| | | Reverse | TTTTAAGCTT**CTGACCTCATGCTAAGCGGCACAC** |
| *ATXN1*-R-PutE2 | chr6:16803384-16804756 | Forward | AAAAGATATC**TAGTGGGCAGTCATTGAAGTAGGG** |
| | | Reverse | TTTTAAGCTT**CCAGAAGAGGAAAGGGAGGGTATT** |
| *ATXN3*-R-PutE1 | chr14:91975120-91975888 | Forward | AAAAGCTAGC**GCCAACTTCTCTGCACCTGTTAACC** |
| | | Reverse | TTTTAGATCT**TGAAGGGCATGTCCCTGCTTGTC** |
| *ATXN3*-R-PutE2 | chr14:92505311-92507475 | Forward | AAAACTCGAG**TGGGCGAAATGGGCGATTATACAC** |
| | | Reverse | TTTTAAGCTT**ACTGGAGGCATTCCCAATGTTCTG** |
| *ATXN3*-R-PutE4 | chr14:93214097-93215784 | Forward | AAAAGCTAGC**TTCACCACATACCACAGGGCTAGTC** |
| | | Reverse | TTTTAGATCT**CCACCCAGAAACACCAACTCCAAAG** |
| *TBP*-R-PutE1 | chr6:170150661-170152979 | Forward | AAAACTCGAG**GGTCCACGAAGTAAGTACGAAGAG** |
| | | Reverse | TTTTGATATC**GGCAGAGGTAAAGTTTAGCCTTCC** |
| *TBP*-R-PutE2 | chr6:170189723-170191350 | Forward | AAAAAAGCTT**CTGCAGAGTGTGACAGTGACATTCC** |

| | | | | |
|---|---|---|---|---|
| | | Reverse | | TTTTGATATC**TAATAAGCCTGTGCTGCTGGCTTCG** |
| *ITPR1*-R-PutE2 | chr3:4344253-4346282 | Forward | | AAAAAAGCTT**AGGGCATAATGAGGCATGTTGGAC** |
| | | Reverse | | TTTTGATATC**AGAAACTCAGGCTCTGGGAAGTAG** |
| *ITPR1*-R-PutE7 | chr3:5019011-5020902 | A | Forward | AAAAGATATC**CCAAGGCGACCTGAAGGAAATAC** |
| | | | Reverse | TTTTAAGCTT**CGGCTTCATCACATGAGTCTCAC** |
| | chr3:5020684-5022750 | B | Forward | AAAAGATATC**TCGCCCACGTTACCAACTTCTCCC** |
| | | | Reverse | TTTTAAGCTT**AGCAGCCTCTGCCGACCTTAACTC** |
| Negative control | chr18:941593-943071 | Forward | | AAAACTCGAG**TGTGAACCACGTGTCCCTCTTAG** |
| | | Reverse | | TTTTGATATC**CCCTGTCATTAAGCGATGCATAGC** |
| Positive control | chr5:89704487-89706712 | Forward | | AAAACTCGAG**TGGAGCCGCAACACTGGAATAAC** |
| | | Reverse | | TTTTGATATC**AGCCCATCCTTCTAGGTCTAACG** |

**Supplementary table 10.** Primers used to amplify PutEs of *ITPR1* for cloning into the pGL4.23 Firefly luciferase reporter vector. Primer sequences are shown in bold and restriction sites in green.

| PutEs | Cloned region (Hg19) | Primers Sequence (5' → 3') | | |
|---|---|---|---|---|
| *ITPR1*-PutE1 | chr3:3840264-3842199 | A | Forward | AAAAGATATC**GCTTTCGCTTAAAGCCGATTCCC** |
| | | | Reverse | TTTTAAGCTT**AGATCCTTCCCAAGGCTCTGACTTC** |
| | chr3:3842041-3843945 | B | Forward | AAAAGATATC**AGCTGGAAAGAGGGCCAAGAGGAG** |
| | | | Reverse | TTTTAAGCTT**GTCGCACAACGCCAAGAGGTGAAC** |
| *ITPR1*-PutE4 | chr3:4762029-4764147 | Forward | | AAAACTCGAG**GGCTTAGGAGATCAGGTGATGAC** |
| | | Reverse | | TTTTGATATC**GGTGAAGCCCTCATTGAACATGTG** |
| *ITPR1*-PutE5 | chr3:4790170-4792499 | A | Forward | AAAACTCGAG**TGCTGGTGTCTTTCTTGTGGGTAG** |
| | | | Reverse | TTTTGATATC**TTTCCGGAGACAGCTCCAAGAATC** |
| | chr3:4792321-4794208 | B | Forward | AAAACTCGAG**GGTGGACTGCGTTGTTGCTATGG** |
| | | | Reverse | TTTTGATATC**GAATATGCAGCACACCGCAGACC** |
| *ITPR1*-PutE6 | chr3:4908637-4910722 | Forward | | AAAACTCGAG**ATACTGCGTGCGGTCATGAAAGGC** |
| | | Reverse | | TTTTGATATC**CCTGACCACAGCAGTAGCTTCATAG** |

**Supplementary table 11.** Primers used to amplify the region overlapping between the CNV deletion and the *ITPR1* 4C-contact peak for cloning into the pGL4.23 Firefly luciferase reporter vector. Primer sequences are shown in bold and restriction sites in green.

| Region of interest | Cloned region (Hg19) | Primers Sequence (5' → 3') | |
|---|---|---|---|
| *ITPR1*-contact peak-deletion | chr3:3837002-3838389 | Forward | AAAACTCGAG**GTCTATGAGCTGGTCACATCATCC** |
| | | Reverse | TTTTGATATC**CGATTGGCTGTGTTGAGACACTAC** |

**Supplementary table 12.** Primers used for Sanger sequencing to confirm the break point of the CNV identified in the patient.

| **Primers** | **Sequence Sequence (5' → 3')** |
|---|---|
| Forward-1 | GAGTAGGGAAGAAACGCACCCAATC |
| Forward-2 | CAAGCTCACTCTGACTGGGTATG |
| Forward-3 | CTGCCCTACCCAATATGGTTGTC |
| Forward-4 | GTGAGCTTGTAAACGGTCCAAGG |
| Forward-5 | TTGCCGACCGGAGAGTCTGAAATC |
| Forward-6 | CACGTGGCCAACTCCCTAACTTC |
| Universal-reverse | TAGGAGGCCAGCTAGGCACAATG |

**Supplementary table 13**. qPCR primers used to measure the expression of *ITPR1* and *GAPDH* and *Beta-actin* in whole blood.

| **Gene** | **Primers Sequence (5' → 3')** | |
|---|---|---|
| *ITPR1*-A | Forward | GTTTCAGCCCTCAGTGGACC |
| | Reverse | CCTTCAGGCACAGAGACCAG |
| *ITPR1*-B | Forward | CCAGTTCAAAAGCCCTGTG |

| | | |
|---|---|---|
| | Reverse | TGTTCCAATACCCTGCTC |
| *GAPDH* | Forward | ATGGGGAAGGTGAAGGTCG |
| | Reverse | GGGGTCATTGATGGCAACAATA |
| *Beta-actin* | Forward | AGCCTCGCCTTTGCCGA |
| | Reverse | GCGCGGCGATATCATCATC |

**Supplementary table 14.** Identified PutEs, R-PutEs and *in vitro*-active enhancers of *ATXN1*, *ATXN3*, *TBP* and *ITPR1*.

| Gene | PutEs | Location (Hg19) | R-PutEs | *In vitro*-active enhancer |
|---|---|---|---|---|
| *ATXN1* | PutE1 | chr6:16319229-16323196 | Yes | Yes |
| | PutE2 | chr6:16803696-16804514 | Yes | Yes |
| *ATXN3* | PutE1 | chr14:91975209-91975777 | Yes | Yes |
| | PutE2 | chr14:92505300-92508216 | Yes | Yes |
| | PutE3 | chr14:93069393-93070523 | No | - |
| | PutE4 | chr14:93214153-93215641 | Yes | No |
| | PutE5 | chr14:93259723-93261534 | No | - |
| *TBP* | PutE1 | chr6:170151018-170153022 | Yes | Yes |
| | PutE2 | chr6:170189867-170191217 | Yes | Yes |
| *ITPR1* | PutE1 | chr3:3839946-3843237 | No | - |
| | PutE2 | chr3:4344258-4346242 | Yes | Yes |
| | PutE3 | chr3:4508178-4509659 | No | - |

| | | | | |
|---|---|---|---|---|
| | PutE4 | chr3:4761716-4763874 | No | - |
| | PutE5 | chr3:4791018-4794112 | No | - |
| | PutE6 | chr3:4908570-4911348 | No | - |
| | PutE7 | chr3:5018875-5022761 | Yes | Yes |

**Supplementary table 15**. Location of the promoters and PutEs of *ATXN1*, *ATXN3*, *TBP* and *ITPR1* with respect to the TADs.

| **Gene** | **Region** | **Location (Hg19)** | **Associated TAD** |
|---|---|---|---|
| *ATXN1* | Promoter | chr6:16759685-16763685 | chr6:16200000-16760000<br>and chr6:16760000-17120000 |
| | PutE1 | chr6:16319229-16323196 | chr6:16200000-16760000 |
| | PutE2 | chr6:16803696-16804514 | chr6:16760000-17120000 |
| *ATXN3* | Promoter | chr14:92570908-92574908 | chr14:91980000-93260000 |
| | PutE1 | chr14:91975209-91975777 | chr14:91980000-93260000 |
| | PutE2 | chr14:92505300-92508216 | chr14:91980000-93260000 |
| | PutE3 | chr14:93069393-93070523 | chr14:91980000-93260000 |
| | PutE4 | chr14:93214153-93215641 | chr14:91980000-93260000 |
| | PutE5 | chr14:93259723-93261534 | chr14:91980000-93260000 |
| *TBP* | Promoter | chr6:170861456-170865456 | chr6:169280000-170900000 |
| | PutE1 | chr6:170151018-170153022 | chr6:169280000-170900000 |
| | PutE2 | chr6:170189867-170191217 | chr6:169280000-170900000 |
| *ITPR1* | Promoter | chr3:4533031-4537031 | chr3:3220000-4680000 |

| | | | |
|---|---|---|---|
| | PutE1 | chr3:3839946-3843237 | chr3:3220000-4680000 |
| | PutE2 | chr3:4344258-4346242 | chr3:3220000-4680000 |
| | PutE3 | chr3:4508178-4509659 | chr3:3220000-4680000 |
| | PutE4 | chr3:4761716-4763874 | chr3:4680000-6840000 |
| | PutE5 | chr3:4791018-4794112 | chr3:4680000-6840000 |
| | PutE6 | chr3:4908570-4911348 | chr3:4680000-6840000 |
| | PutE7 | chr3:5018875-5022761 | chr3:4680000-6840000 |

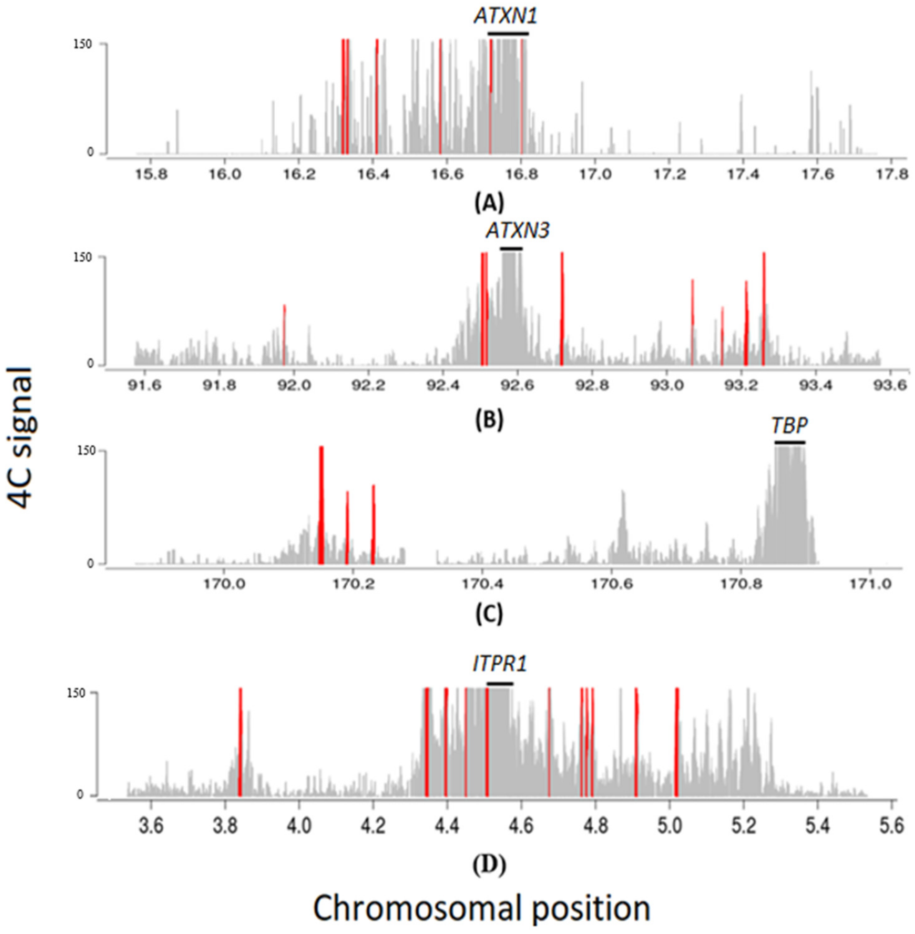


**Supplementary figure 1**. 4C-seq data of *ATXN1*, *ATXN3*, *TBP* and *ITPR1* promoter showing genomic regions significantly interacting with the promoters (4C-contact peaks). The location of the promoter viewpoints is shown as a black line and the 4C-contact peaks as red bars. (A) *ATXN1* promoter has six significant 4C-contact peaks. (B) *ATXN3* promoter has eight significant 4C-contact peaks. (C) *TBP* promoter has three significant 4C-contact peaks. (D) *ITPR1* promoter has eleven significant 4C-contact peaks.

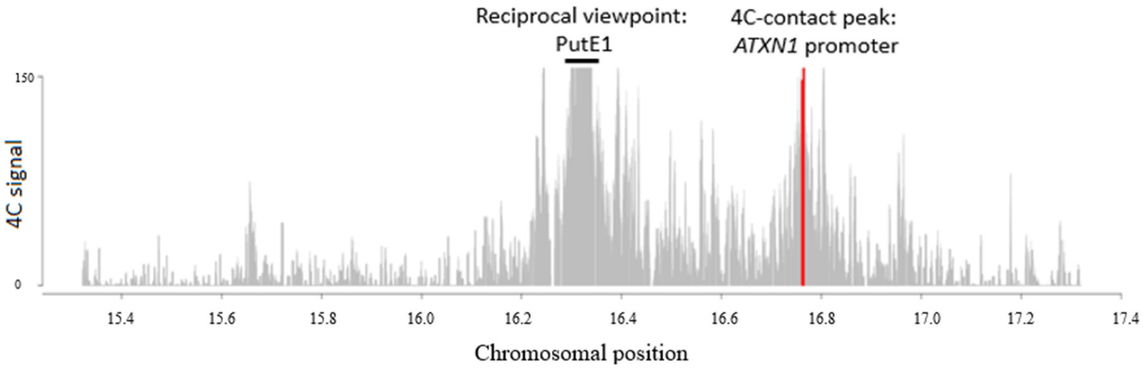


(A)

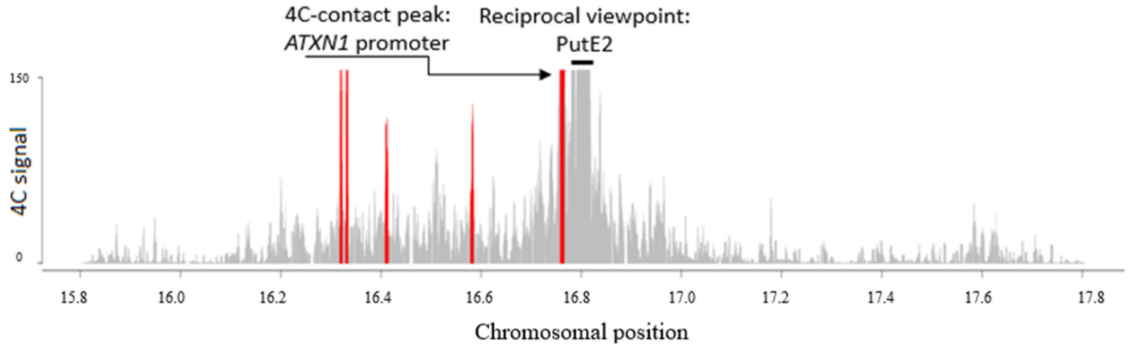


(B)

**Supplementary figure 2**. Reciprocal 4C-seq for the PutEs of *ATXN1*. (A) PutE1 is selected as a reciprocal viewpoint (chr6:16319229-16323196) and *ATXN1* promoter (chr6:16759520-16767014) is a 4C-contact peak for PutE1. (B) PutE2 is selected as a reciprocal viewpoint (chr6:16803696-16804514) and *ATXN1* promoter (chr6:16758209-16766894) is a 4C-contact peak for PutE2. Based on the Eukaryotic Promoter Database, the location of the *ATXN1* promoter is chr6:16759685–1676368. Black lines show the location of the reciprocal viewpoint. Red bars indicate the significant 4C-contact peaks.

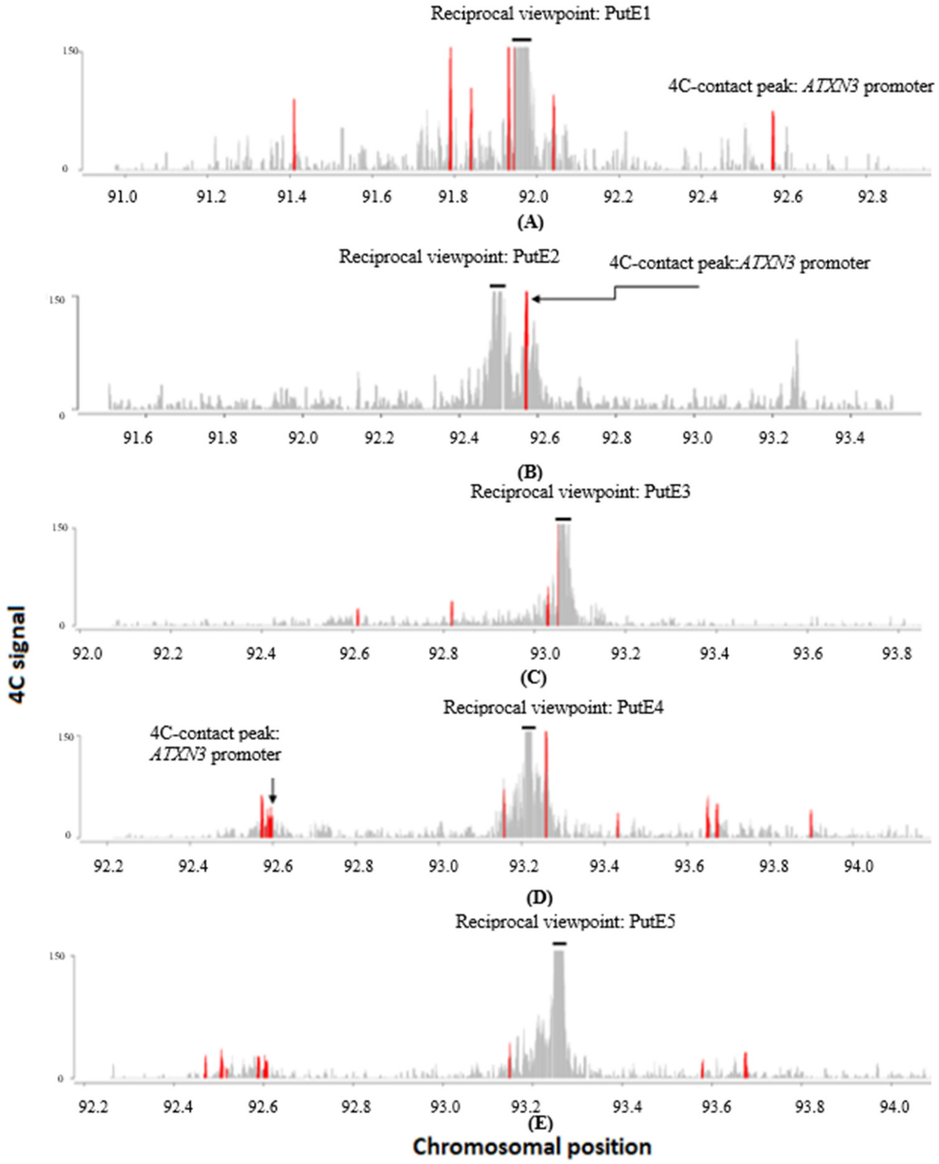


**Supplementary figure 3.** Reciprocal 4C-seq for PutEs of *ATXN3*. (A) PutE1 is selected as a reciprocal viewpoint (chr14:91975209-91975777), and *ATXN3* promoter (chr14:92570837-92577020) is a 4C-contact peak for PutE1. (B) PutE2 is selected as a reciprocal viewpoint (chr14:92505300-92508216), and *ATXN3* promoter (chr14:92567911-92574975) is a 4C-contact peak for PutE2. (C) PutE3 is selected as a reciprocal viewpoint (chr14:93069393-93070523), but *ATXN3* promoter is not a 4C-contact peak for PutE3. (D) PutE4 is selected as a reciprocal viewpoint (chr14:93214153-93215641), and *ATXN3* promoter (chr14:92570462-92599948) is a 4C-contact peak for PutE4. (E) PutE5 is selected as a reciprocal viewpoint (chr14: chr14:93259723-93261534), but *ATXN3* promoter is not a 4C-contact peak for PutE5. Based on the Eukaryotic Promoter Database, the location of the *ATXN3* promoter is chr14:92570908–92574908. Black lines show the location of the reciprocal viewpoint. Red bars indicate the significant 4C-contact peaks.

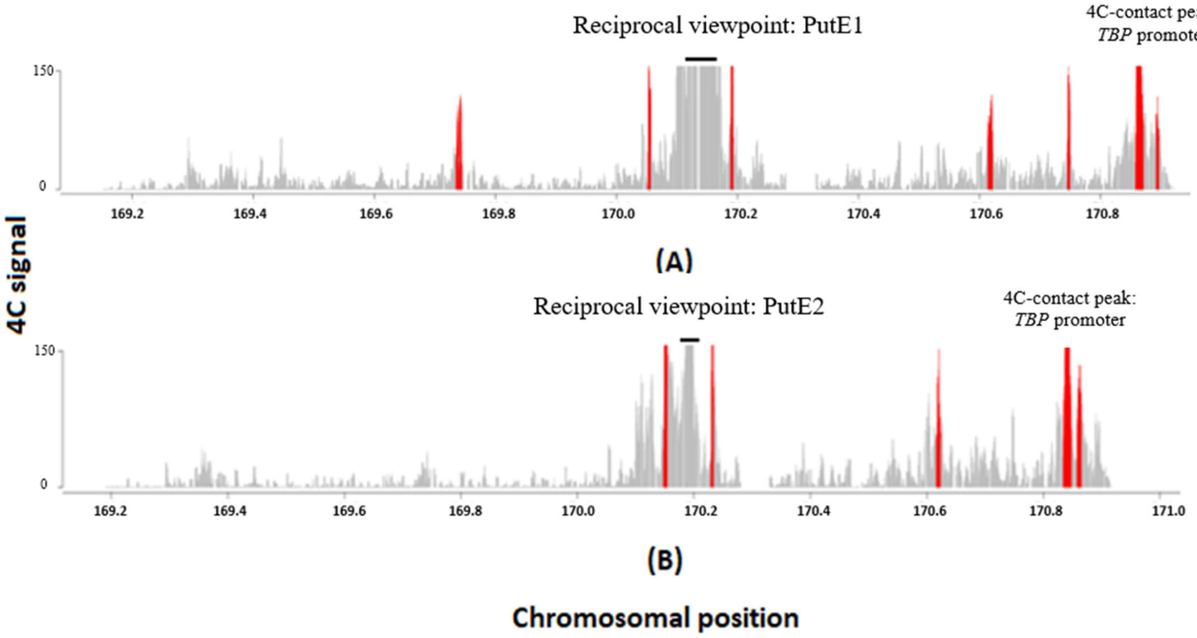


**Supplementary figure 4.** Reciprocal 4C-seq for PutEs of *TBP*. (A) PutE1 is selected as a reciprocal viewpoint (chr6:170151018-170153022), and *TBP* promoter (chr6:170857754-170871249) is a 4C-contact peak for PutE1. (B) PutE2 is selected as a reciprocal viewpoint (chr6:170189867-170191217), and *TBP* promoter (chr6:170857849-170865809) is a 4C-contact peak for PutE2. Based on the Eukaryotic Promoter Database, the location of the *TBP* promoter is chr6:170861456–170865456. Black lines show the location of the reciprocal viewpoint. Red bars indicate the significant 4C-contact peaks.

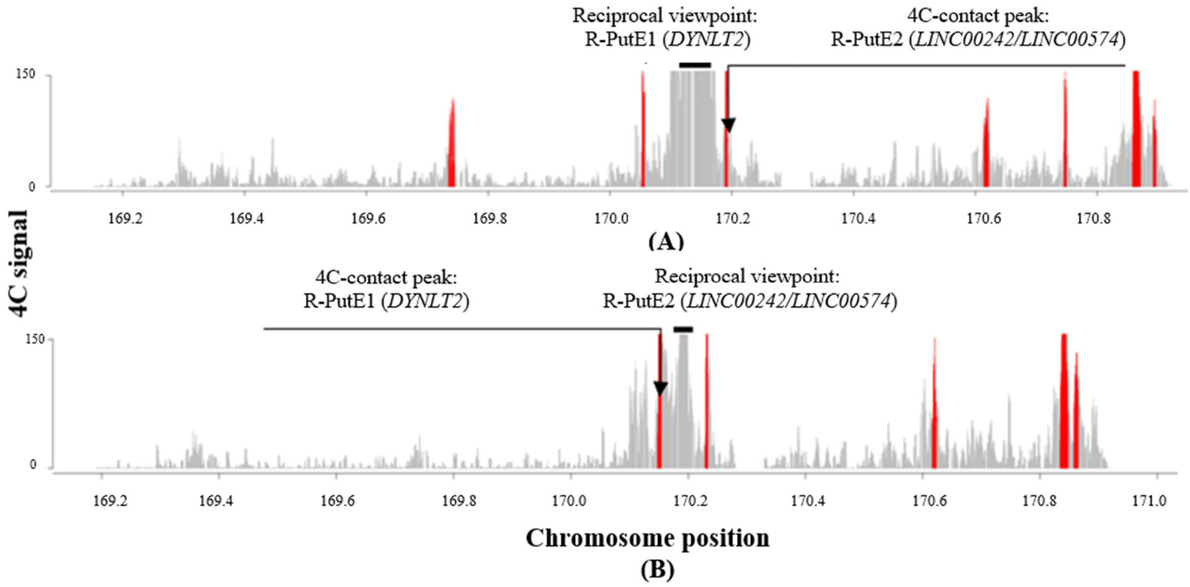


**Supplementary figure 5.** Reciprocal 4C-seq showing an interaction between R-PutE1 and R-PutE2 of *TBP*. (A) R-PutE1, which encompasses *DYNLT2*, is selected as a reciprocal viewpoint (chr6:170151018-170153022) and R-PutE2, which encompasses *LINC00242/LINC00574* (chr6:170188785-170193422) is a 4C-contact peak for R-PutE1. (B) R-PutE2 which encompasses *LINC00242/LINC00574* is selected as a reciprocal viewpoint (chr6:170189867-170191217), and R-PutE1 which encompasses *DYNLT2* (chr6:170147727-170154438) is a 4C-contact peak for R-PutE2. Black lines show the location of the reciprocal viewpoint. Red bars indicate the significant 4C-contact peaks.

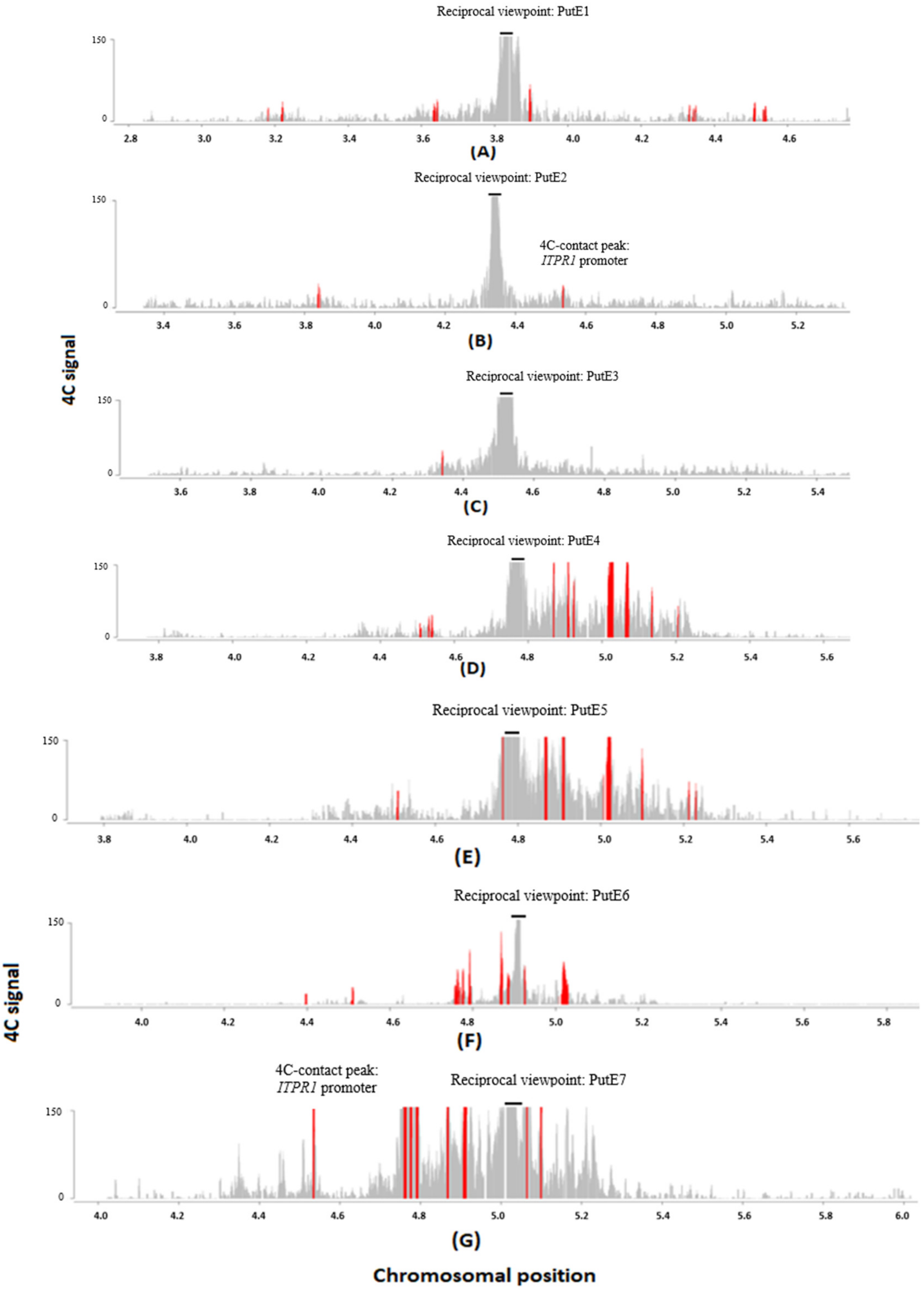


**Supplementary figure 6.** Reciprocal 4C-seq for the PutEs of *ITPR1*. (A) PutE1 is selected as a reciprocal viewpoint (chr3:3839946-3843237), but *ITPR1* promoter is not a 4C-contact peak for PutE1. (B) PutE2 is selected as a reciprocal viewpoint (chr3:4344258-4346242), and *ITPR1* promoter (chr3:4533825-4538002) is a 4C-contact peak for PutE2. (C) PutE3 is selected as a reciprocal viewpoint (chr3:4508178-4509659), but *ITPR1* promoter is not a 4C-contact peak for PutE3. (D) PutE4 is selected as a reciprocal viewpoint (chr3:4761716-

4763874), but *ITPR1* promoter is not a 4C-contact peak for PutE4. (E) PutE5 is selected as a reciprocal viewpoint (chr3:4791018-4794112), but *ITPR1* promoter is not a 4C-contact peak for PutE5. (F) PutE6 is selected as a reciprocal viewpoint (chr3:4908570-4911348), but *ITPR1* promoter is not a 4C-contact peak for PutE6. (G) PutE7 is selected as a reciprocal viewpoint (chr3:5018875-5022761) and *ITPR1* promoter (chr3:4533360-4537206) is a 4C-contact peak for PutE7. Based on the Eukaryotic Promoter Database, the location of the *ITPR1* promoter is chr3:4533031–4537031. Black lines show the location of the reciprocal viewpoint. Red bars indicate the significant 4C-contact peaks.

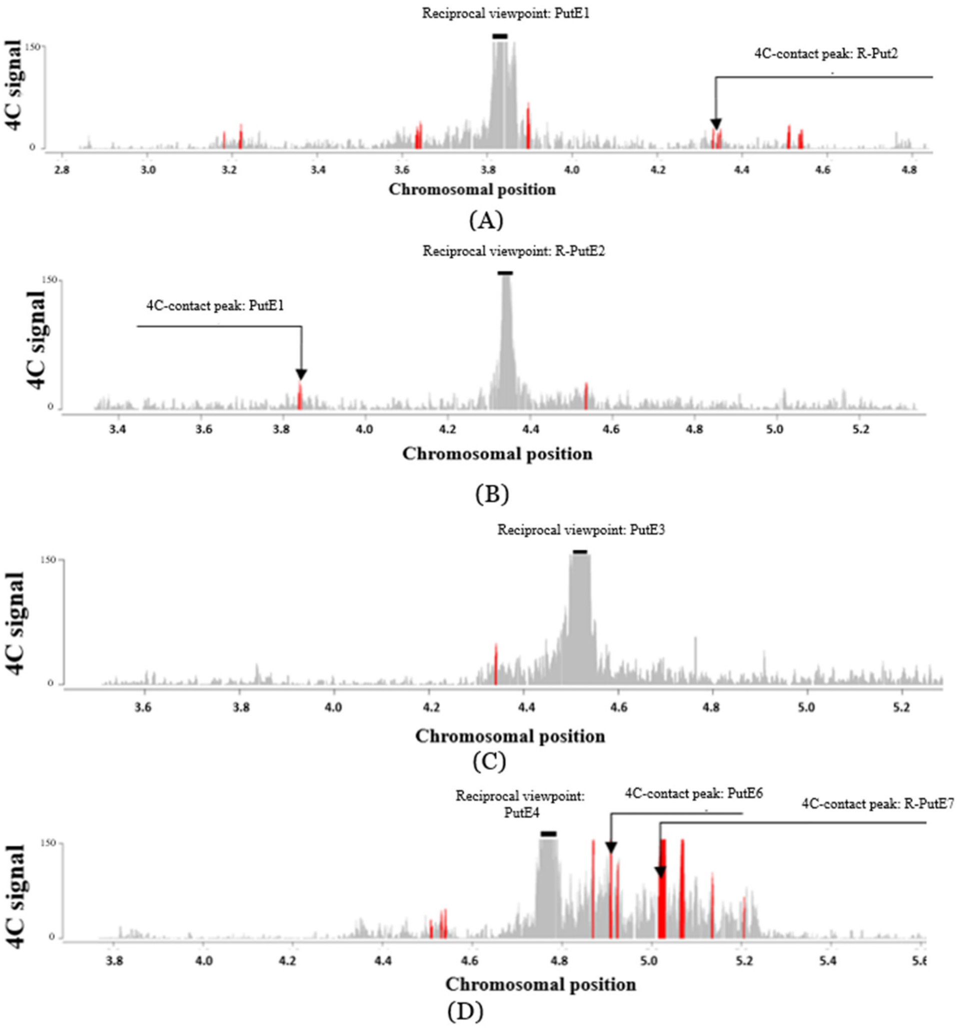

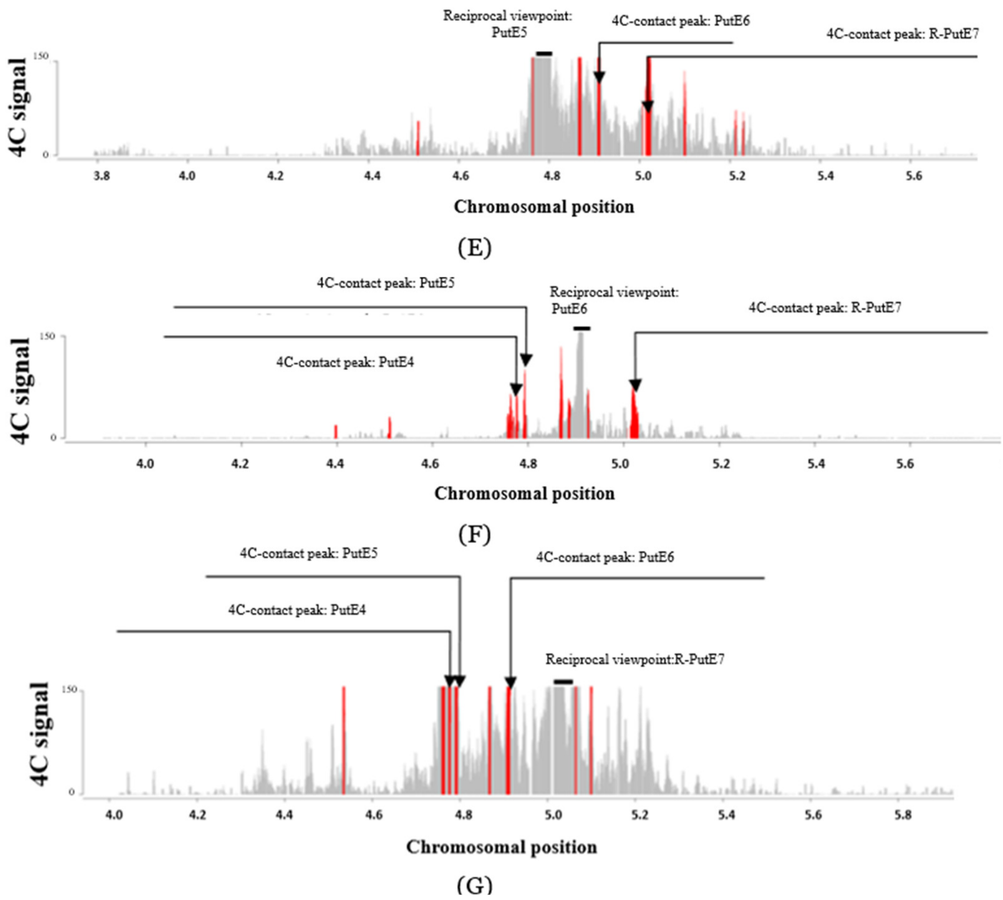


**Supplementary figure 7.** Reciprocal 4C-seq for the PutEs of *ITPR1* showing interactions between the PutEs/R-PutEs. (A) PutE1 interacts with R-PutE2. (B) R-PutE2 interacts with PutE1. (C) PutE3 does not interact with other PutEs/R-PutEs. (D) PutE4 interacts with PutE6 and R-PutE7. (E) PutE5 interacts with PutE6 and R-PutE7. (F) PutE6 interacts with PutE4, PutE5 and R-PutE7. (G) R-PutE7 interacts with PutE4, PutE5 and PutE6.

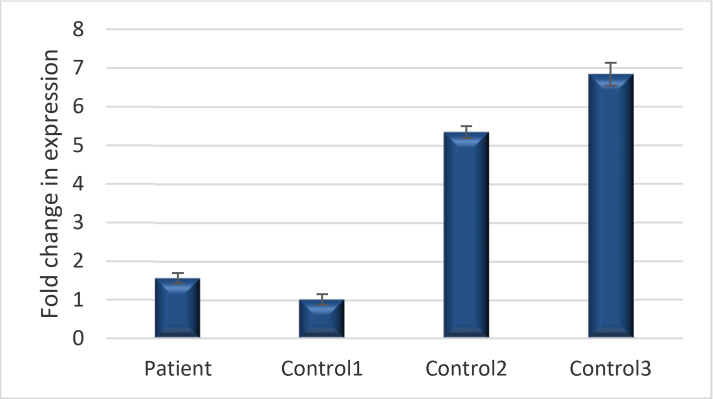


**Supplementary figure 8.** Expression analysis of *ITPR1* in blood from the SCA patient and three healthy individuals using a second set of primers to measure *ITPR1* expression levels for confirmation. The delta-delta Ct method is used to determine the relative fold change in expression levels. Expression value for control 1 is set to 1, with other individuals compared to control 1. The relative fold change in expression for each individual is based on the means of three independent replicates. Standard deviations are shown as error bars.
